# Structure and function of the nervous system in nectophores of the siphonophore *Nanomia bijuga*

**DOI:** 10.1101/2020.07.18.210310

**Authors:** Tigran P. Norekian, Robert W. Meech

## Abstract

Although *Nanomia* nectophores are specialized for locomotion, their cellular elements and complex nerve structures suggest they have multiple subsidiary functions.
The main nerve complex is a nerve ring, an adjacent columnar-shaped matrix plus two associated nerve projections. An upper nerve tract appears to provide a sensory input while a lower nerve tract connects with the rest of the colony.
The nerve cell cluster that gives rise to the lower nerve tract may relay information from the colony stem.
The structure of the extensively innervated “flask cells” located around the bell margin suggests a secretory function. They are ideally placed to release chemical messengers or toxins into the jet of water that leaves the nectophore during each swim.
The numerous nematocytes present on exposed nectophore ridges appear to have an entangling rather than a penetrating role.
Movements of the velum, produced by contraction of the Claus’ muscle system during backwards swimming, can be elicited by electrical stimulation of the surface epithelium even when the major nerve tracts serving the nerve ring have been destroyed (confirming Mackie, 1964).
Epithelial impulses generated by electrical stimulation elicit synaptic potentials in Claus’ muscle fibres. Their amplitude suggests a neural input in the vicinity of the Claus’ muscle system. The synaptic delay is <1.3 ms (Temperature 11.5 to 15° C).
During backward swimming radial muscle fibres in the endoderm contract isometrically providing the Claus’ fibres with a firm foundation.

**Summary Statement:** Nanomia colonies have specialized swimming bells capable of backwards swimming; thrust is redirected by an epithelial signal that leads to muscle contraction via a synaptic rather than an electrotonic event.

## Introduction

The most striking feature of the behaviour of the siphonophore *Nanomia* is that it is able to take evasive action by rapidly swimming either backwards or forwards. In either case movement depends on the contraction of bell-like structures, the nectophores that form a column at the anterior end of the colony (see Figure 1A). Each nectophore resembles a medusa “stripped of its gonads, tentacles, mouth and manubrium” (Mackie et al., 1987). Its ring-shaped nervous system receives an input from the rest of the colony via nerves in the stem-like structure that links the different parts of the colony together. As with other hydromedusan swimming bells the thrust for movement originates in the contraction of a striated muscle sheet, myoepithelium, lining the subumbrella cavity of the bell. Under normal conditions water is forced out of the bell cavity and the bell is jet-propelled forward.

**Figure 1.**
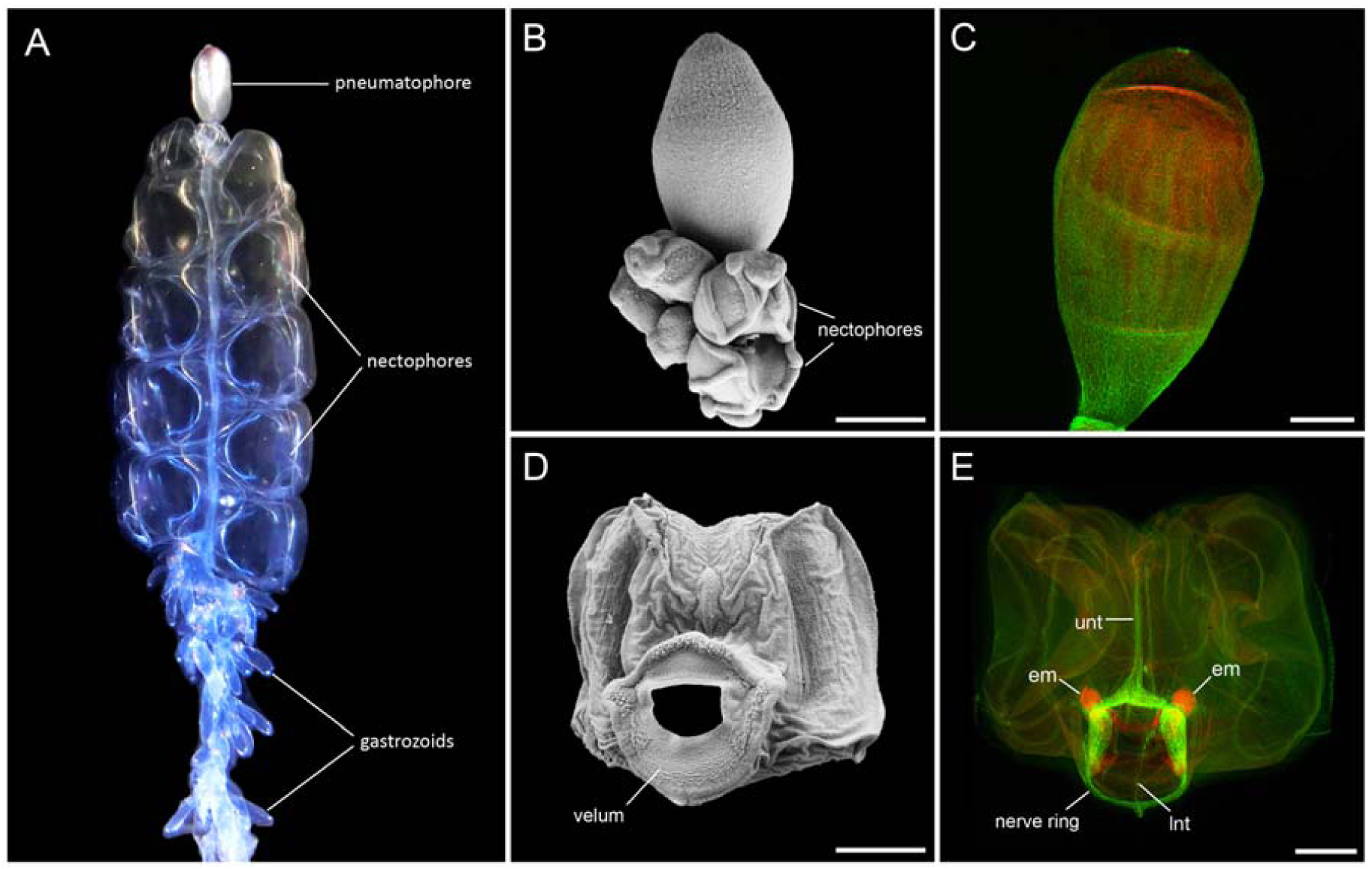
General view of the nectophores of Nanomia bijuga including SEM images (B, C) and antitubulin-IR/phalloidin stained images (D, E). (A) – Anterior region of the colony showing pneumatophore, nectophores and gastrozoids. The stem is visible through the transparent nectophores. (B, D) – Pneumatophore with young nematophores attached at its base. (C, E) – Nematophore. Antitubulin-IR, which labels the neural system, is green, while phalloidin, which labels F-actin is red. Abbreviations: unt – upper nerve tract; lnt – lower nerve tract; em – endodermal muscle fibres. Scale bars: A – 5 mm, B, C, D, E – 500 µm

Mackie (1964) has shown that backward swimming depends on the contraction of specialized radial muscles. These are Claus’ fibres, part of a thin muscular diaphragm (velum) that runs around the bell aperture. When Claus’ fibres contract the water expelled from the bell is redirected forwards.

Given the apparent simplicity of the nectophore nerve circuits, the challenge was to explain the way in which its different muscle groups are controlled. Mackie (1964) found that for forward swimming, signals pass from the stem to the subumbrella musculature via a specific route, the lower nerve tract. On the other hand stimuli that evoke reverse swimming follow no single localized route and are carried all over the outer surface of the nectophore, by way of a non-muscular excitable epithelium. The way in which excitation passed from the excitable epithelium to the Claus’ muscle group was unclear. There was no evidence for the existence of chemical synapses between epithelium and muscle and it seemed possible that the link depended on electrical junctions instead.

This paper presents the neuroanatomy of the nectophore, and shows that although *Nanomia* nectophores are clearly specialized for locomotion, their different cellular elements and nerve structures testify to the existence of other subsidiary functions. Intracellular recordings from Claus’ fibres in the velum show that the action potentials associated with excitation arise from slower, depolarizing events. We suggest that epithelial impulses arriving at the nectophore margin act through ring nerves to excite the Claus’ fibre system synaptically.

## Methods

### Animals

Adult specimens of *Nanomia bijuga* were collected from surface water at the dock of the Friday Harbor Laboratories, University of Washington, USA, and held in 2 l jars of seawater at 7-9°C. Experiments were performed in the spring-summer seasons of 2016-2020.

### Terminology

Haddock et al., (2005) provide a comprehensive terminology to specify the axes of siphonophores. We follow them in using the term “anterior” to describe the end of the colony with the pneumatophore. We also use the terms “upper” and “lower” to replace the terms “dorsal” and “ventral” in isolated nectophores, “dorsal” and “ventral” being retained as descriptors for the entire colony. For the experimental work on isolated nectophores we have found it convenient to use the terms “distal” and “proximal” to the margin to describe muscle locations.

### Scanning Electron Microscopy (SEM)

Nectophores were detached from adult *Nanomia* and fixed in 2.5% glutaraldehyde in 0.1 M phosphate-buffered saline (PBS; pH=7.6) for 3-5 hours at room temperature. They were then washed for several hours in 2.5% sodium bicarbonate. For secondary fixation, the samples remained in 2% osmium tetroxide, 1.25% sodium bicarbonate for 3 hours at room temperature. They were rinsed several times with distilled water, dehydrated in ethanol (10 minutes each in 30, 50, 70, 90 and 100 percent ethanol) and placed in a Samdri-790 unit (Tousimis Research Corporation) for critical point drying. After the drying process, the samples were mounted on holding platforms and processed for metal coating using an SPI Sputter Coater. SEM imaging was performed using a NeoScope JCM-5000 microscope (JEOL Ltd., Tokyo, Japan).

### Immunocytochemistry

Detached *Nanomia* nectophores were fixed overnight in 4% paraformaldehyde in 0.1 M PBS (5° C; pH=7.6) and then washed for 2 hours in PBS. The specimens were pre-incubated for about 6 hours in a blocking solution of 6% goat serum in PBS, and then incubated for a further 48 hours in the primary anti-tubulin antibody diluted in blocking solution at a final dilution of 1:40. The rat monoclonal anti-tubulin antibody (AbD Serotec, Bio-Rad, Cat# MCA77G, RRID: AB_325003) recognizes the alpha subunit of tubulin, and specifically binds tyrosylated tubulin (Wehland & Willingham, 1983; Wehland, Willingham, & Sandoval, 1983). Following a series of PBS washes for 6-8 hours, the specimens were incubated for 24 hours in secondary antibodies – goat anti-rat IgG antibodies (Alexa Fluor 488 conjugated; Molecular Probes, Invitrogen, Waltham, MA, Cat# A11006, RRID: AB_141373) at a final dilution 1:20. To label the muscle fibres, we used the well-known marker phalloidin (Alexa Fluor 568 phalloidin from Molecular Probes, Invitrogen, Waltham, MA, Cat# A11077, RRID: AB_2534121), which binds to F-actin. Following the secondary antibody treatment, the specimens were incubated in phalloidin solution (in 0.1 M PBS) for 8 hours at a final dilution 1:80 and then washed for 6-8 hours in several PBS rinses.

To stain the nuclei, the preparations were mounted in mounting medium with DAPI (Vectashield) on glass microscope slides. The slides were viewed and photographed using a Nikon Research Microscope Eclipse E800 with Epi-fluorescence using standard TRITC and FITC filters and Nikon C1 laser scanning confocal microscope. To test for the specificity of immunostaining either the primary or the secondary antibody was omitted from the procedure. In either case no labeling was detected. This anti-tubulin antibody has been used to label the neural systems in the hydrozoan *Aglantha digitale* (Norekian and Moroz, 2020b) and several ctenophore species (Norekian and Moroz, 2016, 2019a, 2019b, 2020a).

### Intracellular recording

Isolated nectophores stabilized with cactus (*Opuntia*) spines in a transparent Sylgard-coated dish were bathed in seawater at 11-15 °C. They were illuminated from below and 10% isotonic MgCl_2_ used suppress contractions of striated muscle (see Kerfoot et al., 1985). A bipolar stainless-steel stimulating electrode placed on the exumbrella epithelium provided a 1-2 ms stimulating pulse which was adjusted to be just suprathreshold. Contractile responses identified the Claus’ fibre region of the velum. Micropipettes filled with 3 M KCl (resistance 40-50 MΩ) were used to monitor the intracellular response of individual fibres at different sites in the velum. Penetration was accomplished by briefly overcompensating the negative capacitance adjustment on the World Precision Instruments Model KS700 preamplifier. The location of the stimulating electrode and the position of each micropipette were recorded photographically.

Extracellular recording: As before, the locations of the stimulating electrode and the recording pipette were stored photographically but now the recordings were made with a low resistance extracellular glass suction pipette (tip diameter, 10-15 μm) rather than a high resistance intracellular one. The pipette was made from hematocrit glass using a two-step pull. Suction was maintained on the velar surface using an air pump connected to the pipette housing so that contact was retained despite the movement of the Claus’ fibres. Currents were recorded using a custom-made “loose patch” clamp amplifier (Roberts and Almers, 1992).

## Results

### General structure

In essence *Nanomia* consists of a float (pneumatophore) and an extended trailing stem, which runs through the colony’s entire length and serves as an anchor for numerous specialized zooids. Attached to the stem just under the float, are the medusa-like swimming bells (nectophores). Below the nectophores are the gastrozooids, specialized for digestion and with tentacles to trap prey. Also present are transparent bracts, which are thought to contribute to floatation (Jacobs, 1962). Figure 1A is a photograph of the nectophore region of *Nanomia bijuga* showing its relationship to the rest of the colony. Newly developing nectophores arise just below the pneumatophore and an SEM of the region shows several at an early stage of development (Figure 1B), three of them with distinct rounded ridges. Even young nectophores contribute to swimming (Costello, et al., 2015) and all resemble small medusae each with a velar opening facing outwards. The bell undergoes partial collapse during fixation and dehydration, but under SEM the ridge structure is still evident as is the broad velum (Figure 1C).

In Figure 1D the pneumatophore is stained with markers for F-actin, (phalloidin; red), and anti-tubulin IR (green). The anti-tubulin IR stains microtubule inter-repeat regions, present in large numbers in much nervous tissue, and the entire surface appears covered by a network of fine nerves supporting the view that it has an important sensory function (Church et al., 2015). In Figure 1E the nectophore is orientated to match Figure 1C and the anti-tubulin staining shows the main nerves; the upper nerve tract (unt), the lower nerve tract (lnt) and the nerve ring. Phalloidin staining reveals the position of an important endodermal muscle (em).

Two sets of muscle fibres are of particular importance for the current investigation. They are the Claus’ fibres, located symmetrically on the upper right and upper left of the velum aperture, and the pair of endodermal fibres that abut them. These muscle groups are brought into play during backward swimming. In Figure 2A the velum is shown with arrows to indicate the position of Claus’ fibres. Fixation has caused the Claus’ fibres to contract and the image approximates to the form that the velum takes during backward swimming. In Figure 2B (same orientation as Figure 2A) the Claus’ fibres are revealed by phalloidin staining (red). Phalloidin also stains other velar muscles as well as the endodermal muscles. The nerve ring (green) encircles the nectophore at the edge of the velum. It consists of an “inner” and an “outer” ring divided by a 0.5 μm thick mesogloea (Jha and Mackie, 1967) but the separation is not visible at this magnification. The upper nerve tract, enters the nerve ring at the top of the figure, while the lower nerve tract, which arises from the back of the nectophore, enters at the bottom.

**Figure 2.**
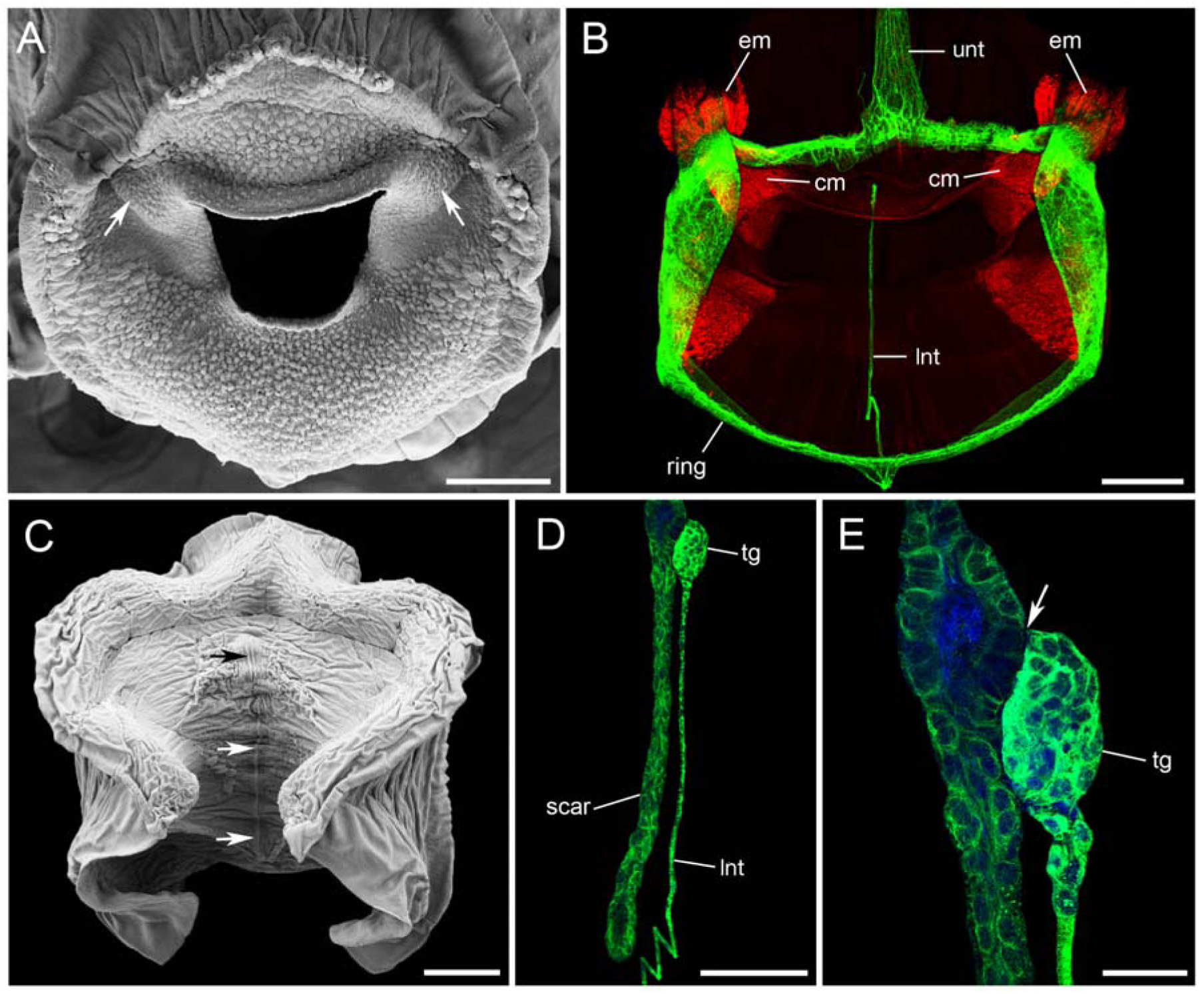
Nectophore structure and innervation. (A) – Velum, SEM image; arrows show location of Claus’ muscle fibres; contraction caused by glutaraldehyde fixation. (B) – Nerve elements labeled with anti-tubulin IR antibody (green); F-actin in muscles labeled with phalloidin (red). (C) – Nectophore from the back, SEM image; arrows show the connection to the stem (scar). (D) – Lower nerve tract labeled with anti-tubulin IR antibody (green), ends in a small “terminal group” of cells attached to the center of the scar. (E) – Arrow shows the point of attachment between the “terminal group” and the epithelial scar. Abbreviations: cm – Claus’ muscle fibres; em – endodermal muscles; unt – upper nerve tract; lnt – lower nerve tract; tg – terminal group. Scale bars: A, B – 200 µm. C – 400 µm; D – 150 µm; E – 40 µm.

### Lower nerve tract

The lower nerve tract, which travels in the ectoderm and enters the nerve ring from the under surface of the nectophore, can be traced backwards to the point of contact between the nectophore and the stem. The whole pathway is seen in Figure 2B, although, because the nectophore has been compressed slightly for the purposes of photography, its passage is “kinked” by a fold in the nectophore wall. At the back of each isolated nectophore there is a narrow scar, visible both in SEM (arrows in Figure 2C) and under confocal microscopy (Figure 2D). This we take to be the point of separation between nectophore and stem (autotomy point). The lower nerve runs to the middle of the scar and ends in a small group of closely attached, anti-tubulin IR staining cells. This “terminal group” (Figure 2D) consists of 40-50 tightly packed cells each about 6-8 µm in diameter and each with a single large nucleus (Figure 2E). The lower nerve tract arises from within the terminal group and does not branch until it arrives at the nerve ring. Its function is to initiate forward swimming by carrying impulses from the stem (Mackie 1964).

### Upper nerve tract

In contrast to the lower nerve tract, the upper nerve tract is highly branched and covers a large area of the top and back surfaces of the nectophore. It connects to the nerve ring with a wide delta-shaped base (Figure 3A). On either side of the delta, running parallel to the main tract, many short neurites merge with the nerve ring (arrows in Figure 3A). These short neurites extend outwards to the exumbrella epithelium. The delta area itself is a complex structure containing numerous nuclei and resembles a ganglion more than a simple collection of nerve fibres (Figure 3B). Although regions in the delta area and main tract sometimes show phalloidin staining (see also Grimmelikhuijzen et al., 1986) the presence of co-localized anti-tubulin IR labeling argues against their being muscle units and we take them to be neurites with unusually high levels of F-actin.

**Figure 3.**
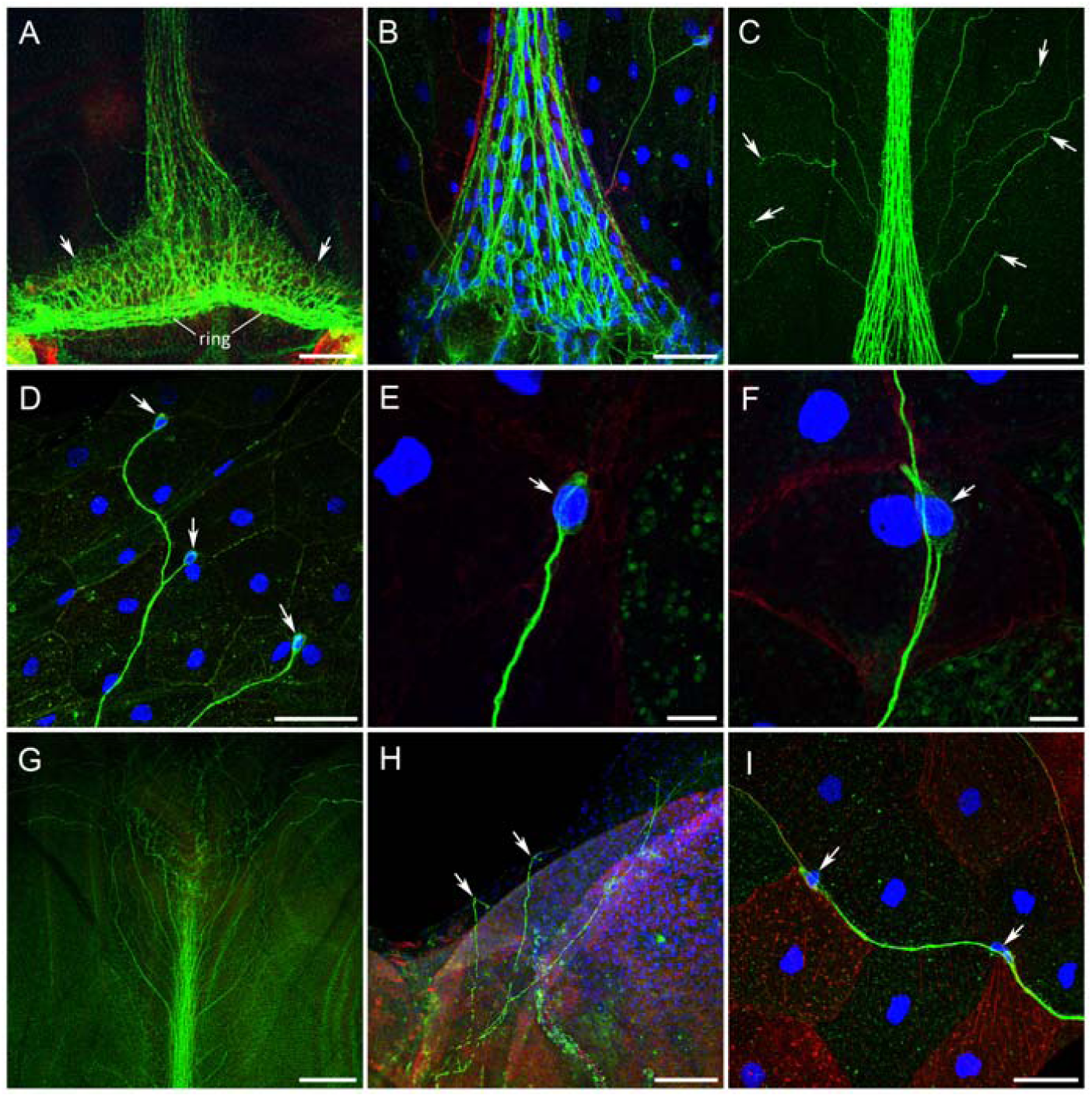
Upper nerve tract. stained with anti-tubulin (green), phalloidin (red) and DAPI (blue). (A) – The delta region at the base of the upper nerve tract where it connects to the nerve ring (ring). Note the many short neurites that connect to the nerve ring (arrows). (B) – Delta region at higher magnification showing numerous nucleated cells. Note phalloidin labeling within the tract itself. (C) – Proximal end of the upper nerve tract near its base showing dispersed branches and oval cell bodies (arrows). (D) – Cell bodies (arrows) of unipolar neurons whose branches travel together towards the main trunk of the upper nerve. (E) – Neural cell body at higher magnification with a large DAPI-stained nucleus (arrow). (F) – Unipolar neuron (arrow) alongside a nerve branch travelling towards the upper nerve tract. (G) – Branches of the upper nerve tract that cover the back of the nectophore. (H) – Nerve branches in the epithelial layer of the exumbrella (arrows); the phalloidin-stained striated muscle layer in the subumbrella is visible through the mesogloea. (I) – A nerve branch navigating its way between large epithelial cells outlined by phalloidin and tubulin-IR staining, 4 µm thick section. Arrows show neural cell bodies. Scale bars: A – 100 µm; B – 40 µm; C – 200 µm; D – 50 µm; E, F – 10 µm; G – 200 µm; H – 100 µm; I – 40 µm.

Beyond the delta area, at the top of each nectophore, the upper tract is joined by numerous thin lateral neurites (Figure 3C). At its distal end the tract breaks up into a mass of fine neurites that cover the entire back surface of the nectophore (Figure 3G). Most branches end in neural cell bodies, each having a single DAPI-stained nucleus (Figure 3C, D). The cell bodies are found at dispersed locations and are slightly elongated spheres 8-10 µm in diameter (Figure 3E, F). They are also located mid-way along neural fibers (Figure 3F, I) and their processes add to the common neural thread (Figure 3F). We suggest that these cells are sensory in nature, gathering information from all over the back of the nectophore, their processes joining to form thicker and thicker nerves until they reach the main upper nerve tract. In much the way a river collects water from numerous small streams, information apparently flows from the periphery to the nerve ring by way of the main upper nerve tract.

All branches of the upper nerve tract run in the exumbrella epithelium (Figure 3H) navigating their way between the epithelial cells. Figure 3I shows a thin branch of the upper nerve tract running precisely between the epithelial cells, clearly outlined by phalloidin and anti-tubulin IR staining (optical section less than 4 µm thick).

### Nerve ring

In common with other hydrozoan swimming bells (Satterlie and Spencer, 1983), *Nanomia* nectophores have a double nerve ring at the edge of the velum which acts as a command center responsible for swimming (Figure 2B). In optical serial sections the nerve ring can be seen to dive downwards towards the junction between the Claus’ and endodermal muscle groups (see Figure 4C, arrow).

**Figure 4.**
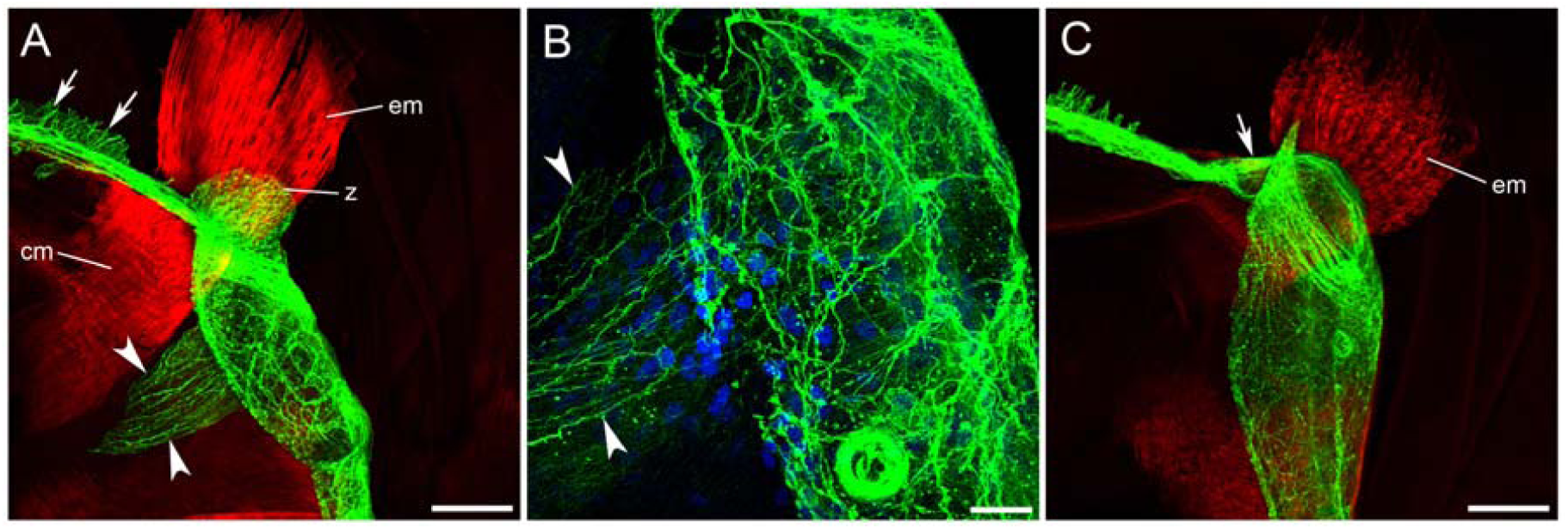
Nerve ring and associated neural network. (A) – Nerve ring crossing the junction between Claus’ muscle fibres (cm) and endodermal muscle fibres (em). A dense network of neural fibres and cell bodies is attached to the nerve ring on its lateral side next to the Claus’ muscle group. Two projections of neural fibers extend from each network. One projection (z) extends toward the “seitliche Zapfen” area (Claus, 1878) in the epithelium. The second is a cone of neural fibers (arrowheads) that projects into the velum toward the Claus’ muscles. Arrows show the short neural processes on the upper side of the nerve ring next to the upper nerve tract, which extend to the epithelial layer. (B) – The dense neural network at high magnification. Arrowheads show the cone shaped projection. (C) – As (A) but from a slightly different angle showing how the ring nerve bends and dives toward the junction between Claus’ fibres and the endodermal muscle. Scale bars: A, C – 100 µm; B – 25 µm.

The nerve ring is not the only nerve complex present in the nectophore. Figure 4 shows that near the Claus’ fibre region, and connected with the nerve ring, is a dense network of neural cell bodies and fibres that form a wide somewhat cylindrical structure. Two of these lateral networks, located symmetrically on either side of the nectophore opening, appear to lie beneath a region containing light sensitive chromatophores (Mackie, 1962). Each lateral network acts as a base for two projections of neural fibers that extend on either side. One projection, the “z” projection, is shorter and directed outwards towards the bell surface (Figure 4A). It appears to innervate the area called the “seitliche Zapfen” (Claus, 1878) – an epithelial thickening that acts as a landmark on either side of the exumbrella. The second projection is a cone-shaped fringe of long nerve processes, extending towards the velum and Claus’ muscle group (Figure 4A, B, arrowheads). Nearby, in each of the upper corners of the velum, are groups of large flask-shaped structures that stain with anti-tubulin antibody (Figure 5A, B). Similar rounded cells are found in another densely innervated area at the base of the upper nerve tract where it joins the ring nerve (Figure 5A). SEM imaging suggests that these cells are secretory in nature (Figure 5C, D). Many of them appear to have released their contents making their openings evident (arrows).

**Figure 5.**
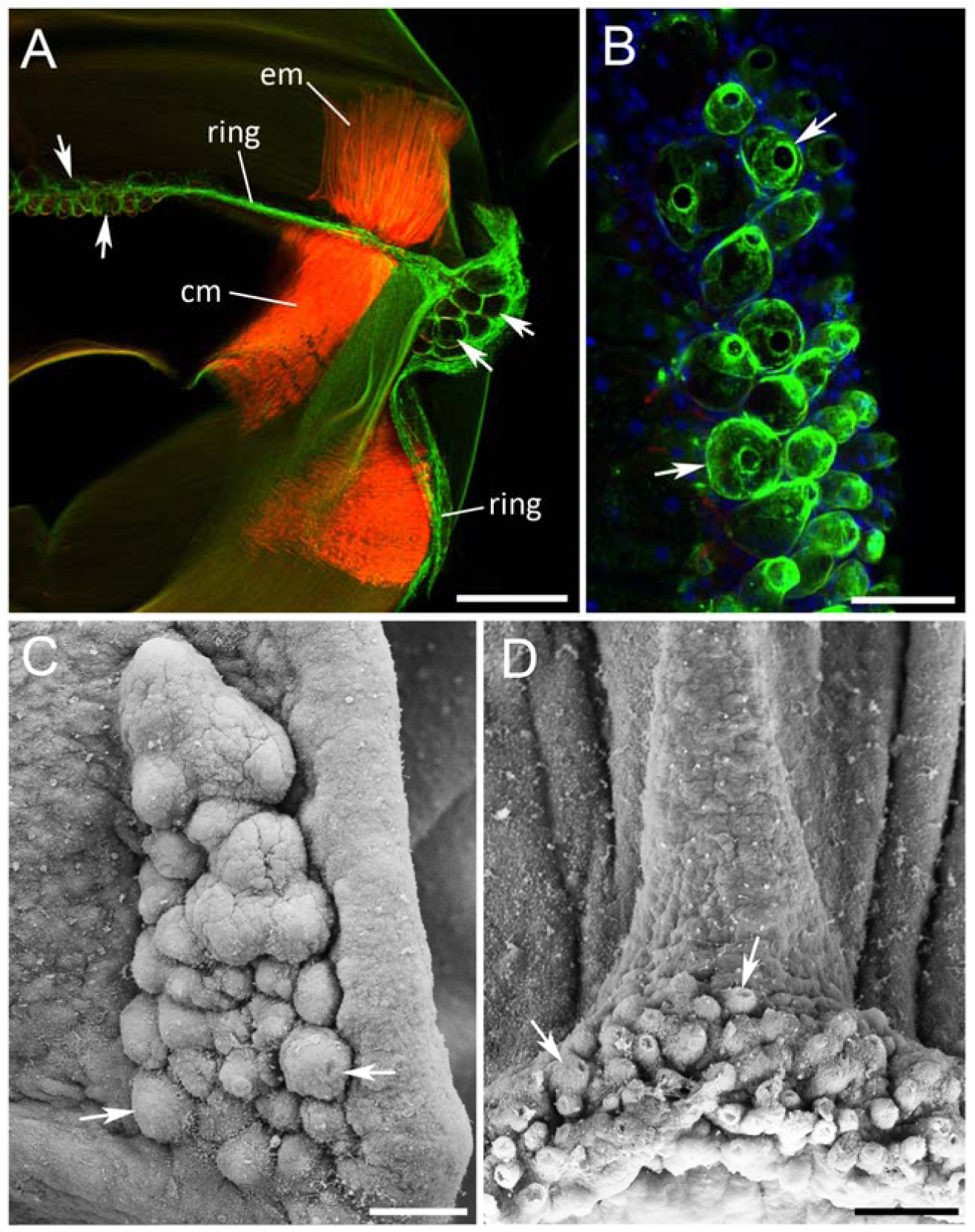
Secretory cells in the nectophore velum. (A) – Anti-tubulin antibody (green) labels two groups of secretory cells located along the nerve ring. One, central, group is at the base of the upper nerve tract (arrows at left). The second, lateral, group is exactly above the symmetrical nerve networks on either side of the velum (arrows at right), where Claus’ muscle fibres (cm) meet the endodermal muscle (em). (B) – Higher magnification of lateral secretory cells labeled by anti-tubulin antibody. (C) – SEM image of lateral secretory cells. (D) – SEM image of central secretory cells. Scale bars: A – 200 µm; B, C, D – 50 µm

### Nematocytes

*Nanomia* nectophores have a distinctive pattern of exumbrellar ridges and depressions that reflect the way they pack together along the stem. These ridges, particularly the two on the upper surface that face the outside, are populated by rows of nematocytes. These are seen as anti-tubulin positive, cone-shaped cells (Figure 6A-C) having a single short cilium (arrows in Figure 6C, D) and a single large nucleus (Figure 6D, E). At the base of each cilium is a narrow ring that stains with phalloidin (Figure 6C-E). Branches of the upper nerve tract frequently cross or run along the ridges (Figure 6D), and sometimes overlap with individual nematocytes (Figure 6E).

**Figure 6.**
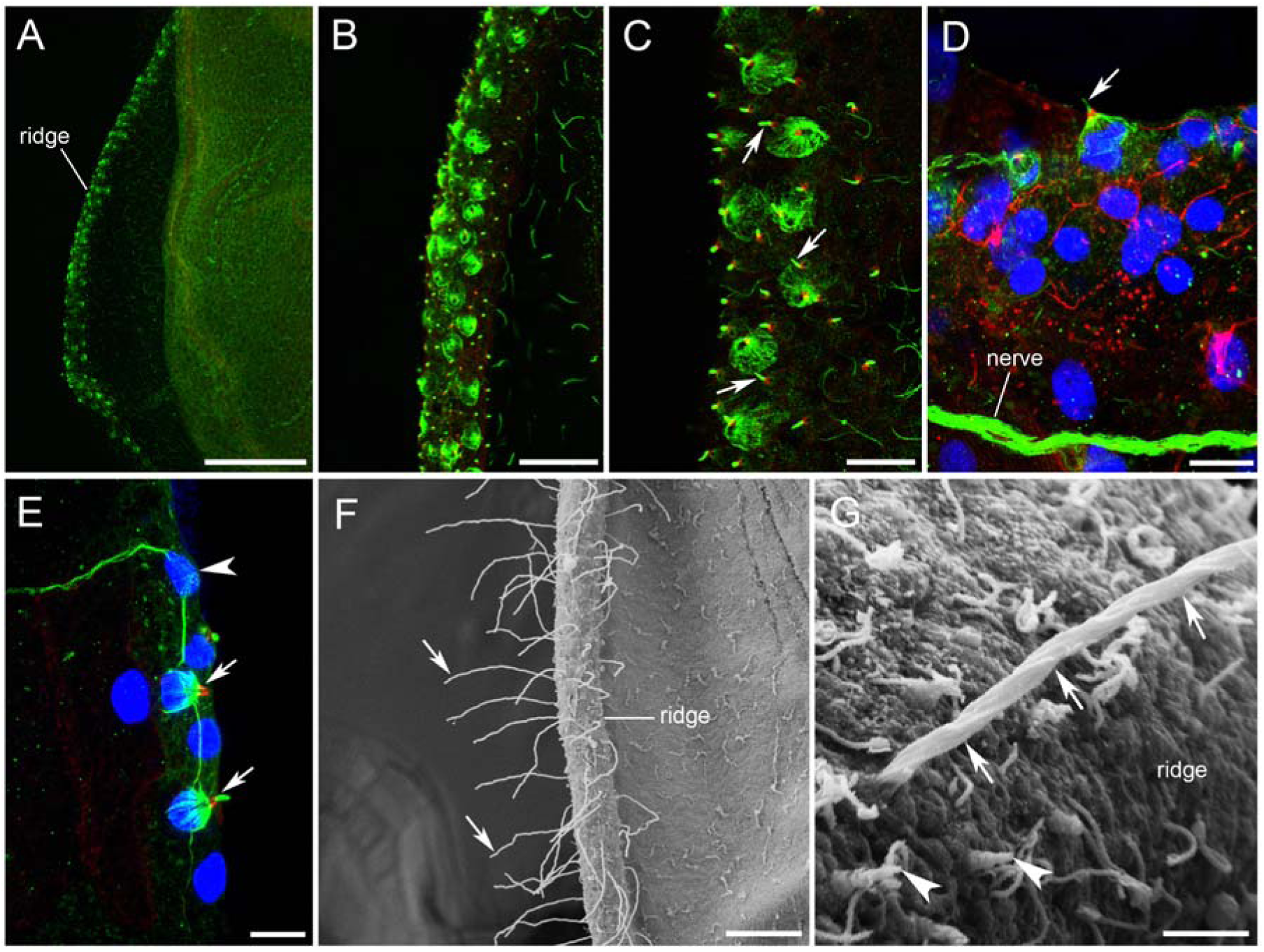
Nematocytes in nectophore ridges. stained with anti-tubulin antibody (green), phalloidin (red) and DAPI (blue). (A, B) – Nectophore ridges with numerous cone-shaped nematocytes. (C) – Nematocysts with single short cilium (cnidocil; arrows). (D) – Base of cilium and outline of individual exumbrella epithelium cells, labeled with phalloidin. (E) – Fine neural process overlapping two nematocytes (arrows). Arrowhead shows bipolar neural cell body, which sends a process to the upper nerve tract. (F) – SEM image showing hair-like structures along the nectophore ridges (arrows). (G) – At higher magnification hairs consist of individual filaments tightly twisted together (arrows). The tubule is undifferentiated along its length and has three rows of serrations projecting about 0.25 μm. Arrowheads show short protruding cilia, which may be the cnidocils of undischarged nematocytes. Scale bars: A – 200 µm; B, F – 50 µm; C, D – 25 µm; E – 10 µm; G – 5 µm.

In many SEM images nectophore ridges exhibit rows of hair-like structures, 70-80 µm in length and 1.5 µm thick, which we take to be tubules of discharged nematocysts (Figure 6F, G) triggered by the glutaraldehyde fixation process. Untriggered nematocysts can be identified by a short sensory cilium – the cnidocil (Figure 6G). At higher magnification the discharged tubules can be seen to consist of several helical filaments tightly twisted together (Figure 6 G). Three filaments have serrated edges. Tubules are non-tapering, with square ends and are closed at the tip. They correspond to the astomocnidae class of nematocyst (Weill, 1934), appear to be non-penetrating and are thought to entangle their targets (Mariscal, 1974; Ostman, 2000).

### Muscle structure

The redirection of the swimming thrust in *Nanomia* is achieved through the contraction of two symmetrical sets of paired radial smooth muscles. Each pair consists of Claus’ fibres in the velum abutting a second muscle group in the endoderm (Figure 2B), their combined action representing a truly remarkable specialization. At high magnification the junction between the two sets of fibres shows a clear separation (Figure 7A) with the ends of endodermal fibres being attached to what appears to be a basement membrane (Figure 7C, arrows). Each group of Claus’ fibres is about 200 µm wide, 300 µm long, with individual muscle “tails” being 3-5 µm thick. Claus’ fibres are myoepithelial cells, like other cnidarian muscles, and their epithelial cell bodies line the outer surface of the velum. The distal ends of the endodermal muscle fibres are in contact with the endodermal lamella (Figure 7B), an epithelial layer attached to the thin mesogloea that lies above the swim muscles of the subumbrella. This circular muscle layer is about 10 µm thick (fibre diameter; 2 µm) and shows clear striations with phalloidin labeling (Figure 7D).

**Figure 7.**
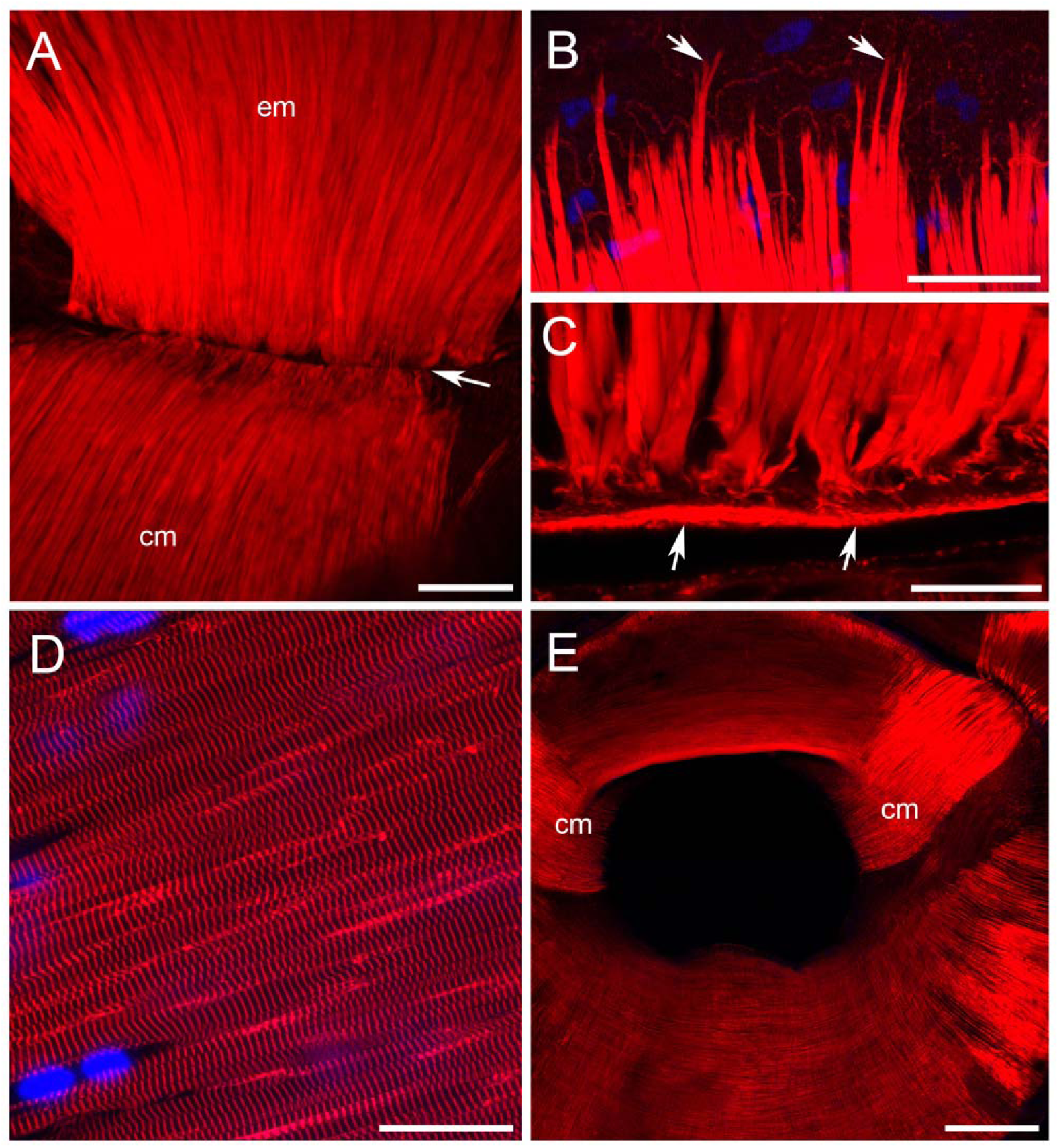
Nectophore muscles. labeled by phalloidin. (A) – Junction (arrow) between Claus’ muscle fibres (cm) and the endodermal muscle fibre group (em). (B) – Distal ends of endodermal muscle fibres terminate in an endodermal epithelium attached to the mesogloea. (C) – Proximal ends of endodermal muscle fibres attached to a basement membrane (arrows) at the point of contact with Claus’ muscle group. (D) – Striated circular muscle fibres in the subumbrella ectoderm. (E) – Smooth radial muscle fibres on the exumbrella surface and striated circular muscle fibres on the subumbrella surface of the velum. Scale bars: A – 50 µm; B, C, D – 25 µm; E – 200 µm.

In addition to the Claus’ fibres, the velum also has other, less well-developed radial smooth muscle fibers on its exumbrella surface. Its subumbrella surface is entirely covered by circular muscle fibres, which are striated under high magnification. In other hydromedusae these two muscle groups control the shape of the velum during swimming (Gladfelter, 1972; Figure 7E).

### Claus’ fibre movements

Photographic images collected at 30 frames/second (see Figure 8A) show that stimulation of the exumbrella produces only a relatively small change in the velum. The main effect is a slight widening at the upper edge of the velum aperture. Other velum muscles remain relaxed so that contraction of the Claus’ fibres is just sufficient to deflect the expelled water onto the concave surface formed by the ballooning of the velum’s lower half (see Figure 4 C in Mackie, 1964).

**Figure 8.**
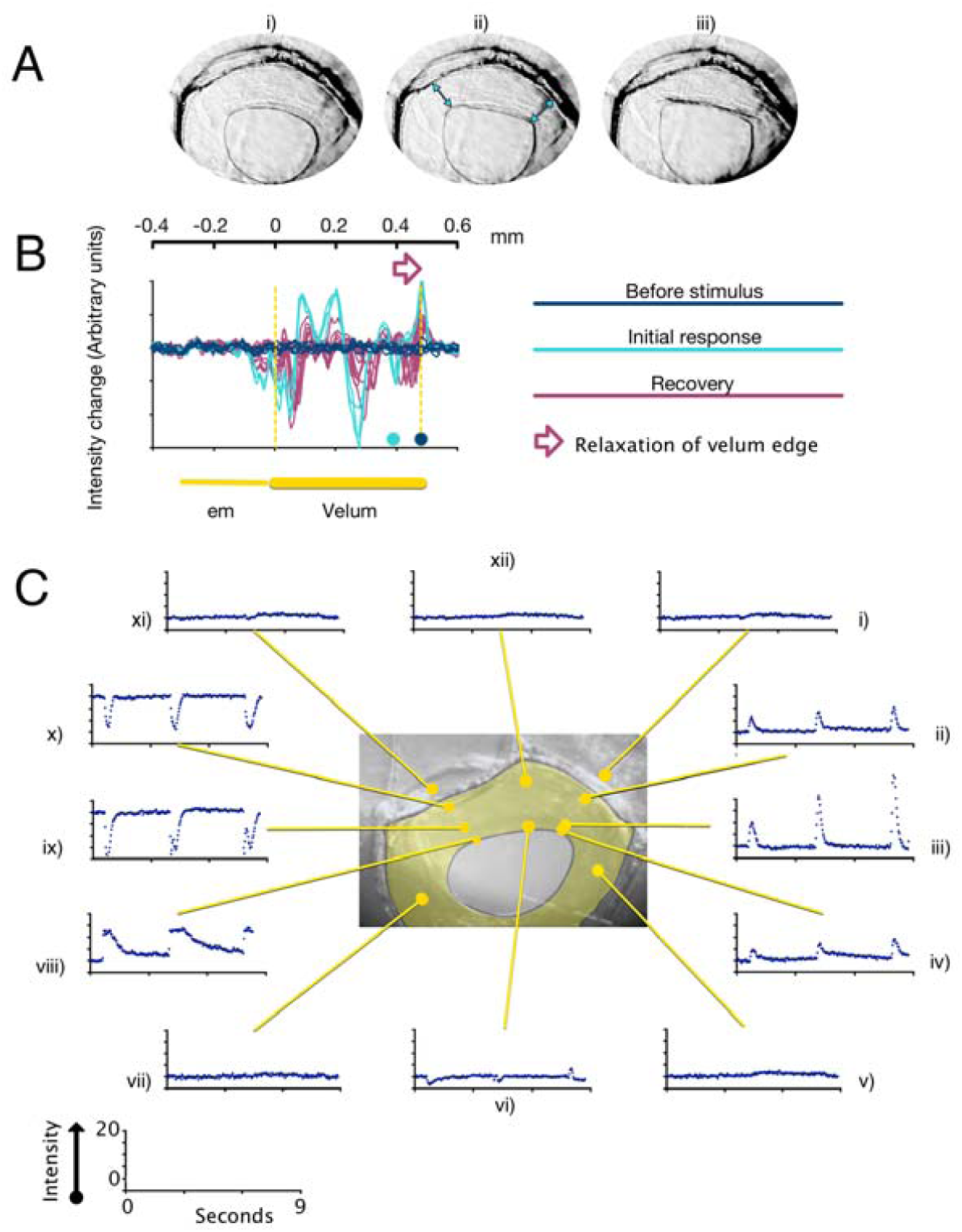
Velum movement with epithelial stimulation. A – Images collected at 30 frames/s before (i) and after (ii, iii) epithelial stimulation. Arrows in ii) indicate the position of the Claus’ fibres. B – changes in transmitted light intensity measured pixel by pixel as described in the text. Black data line, change from baseline before stimulus; blue data line, change from baseline for the three frames immediately after stimulus; purple line, change from baseline during recovery. The nectophore bell margin is at 0 mm. The extent of the velum before epithelial stimulation is shown by the yellow dotted lines. Distal edge of the velum indicated by filled circle before (black) and after (blue) epithelial stimulation. C – time course of the average change in light intensity associated with a series of three suprathreshold stimuli at different points on the velum surface. A, B, C, three different isolated nectophores bathed in normal seawater with 20 mM Tris-HCl pH 7.6. Bipolar stimulating electrode located on left hand side of the exumbrella epithelium, 1.4 mm from the velum margin. Temperature, 13°.

Claus’ fibres are restricted to a narrow strip on either side of the upper velum but their contraction (arrows in Figure 8Aii) stretches the entire mid-region (below the delta area), producing a marked fold just within the velum’s distal edge (Figure 8Aiii). To examine the time course of the Claus’ fibre contraction, video sequences were analyzed frame by frame. Figure 8B shows changes in a typical series of transects crossing a contractile region after epithelial stimulation. In each transect the change in transmitted light intensity is plotted pixel by pixel. To provide a baseline, the transmitted light intensity was recorded over a 500 ms period (i.e. 15 frames) with the nectophore at rest, and an average pixel intensity calculated. The position of the transect remaining constant, the baseline was subtracted from intensity data derived from subsequent video frames. The series shown in Figure 8B consists of a further seven frames collected before the stimulus (black traces; 67 ms interval), three frames following the stimulus (blue traces; 67 ms intervals) and thirteen frames collected during recovery (purple traces; five at 67 ms intervals followed by eight at 334 ms).

From Figure 8B it is evident that a) the effects of muscle contraction are largely confined to the velum region (between dotted yellow lines) and b) radial movement is confined to the distal rim of the velum (open arrow). The solid black circle on the lower axis of the chart shows the position of the distal rim of the velum at rest. At this point the immediate effect of the stimulus is an increase in intensity as a consequence of the retraction of the velum. Over the course of the video sequence the pixel intensity slowly returns towards its resting level.

At the point indicated by the solid blue circle the immediate effect of the stimulus is to reduce pixel intensity, the peak of this reduction corresponding to the edge of the velum after contraction. During the recovery sequence, as the distal rim of the velum gradually relaxes back into its resting position, the peak reduction in transmitted light moves in the distal direction (shown by open arrow).

Not all the intensity changes in Figure 8B arise from radial displacements of the distal velum. In other regions intensity changes follow from the movement of the distal velum as it pushes more proximal regions into ridges and troughs. These folds appear immediately after the stimulus and then subside during the course of recovery without greatly changing their position. Apart from in the vicinity of the nerve ring (yellow dotted line at 0 mm) there is little intensity change of any kind outside the velum. This includes in the region overlying the endodermal muscle fibres.

Figure 8C shows the effect of repeated stimuli on different positions in and around the velum. There are marked changes in average transmitted light in Claus’ fibre areas, but minimal change at other sites. This is true not only in the lateral areas but also in the region at the top of the velum immediately under the delta region. More distally in this mid-region, small changes are evident caused by contractions in neighbouring Claus’ fibres.

In these experiments the stimulating electrode was positioned on the far left hand side of the nectophore (outside of the field of view) and the responses from its nearest Claus’ fibres were slightly stronger than from those on the right. To test whether a small delay might be responsible for the imbalance between the two sides, a cut was made in the mid region of the velum. As might be expected, once this was done the movements caused by Claus’ fibre contractions were greatly enhanced. Movements in the area above the endodermal muscle fibres were also seen. We suppose that the conformation of the velum during backwards swimming depends on a delicate balance between Claus’ fibre contractions, endodermal muscle contractions and the tension in the sub-delta region.

### Electrophysiology

#### Intracellular recording

the effect of stimulating the exumbrella on the Claus’ fibre system was examined using intracellular micropipettes. Figure 9A shows a schematic representation of the velum, traced from stored images, for five nectophore preparations. In the figure the velum is yellow and the dashed green lines indicate the approximate limits to the visible contractions (see Figure 8A for a photographic representation).

**Figure 9.**
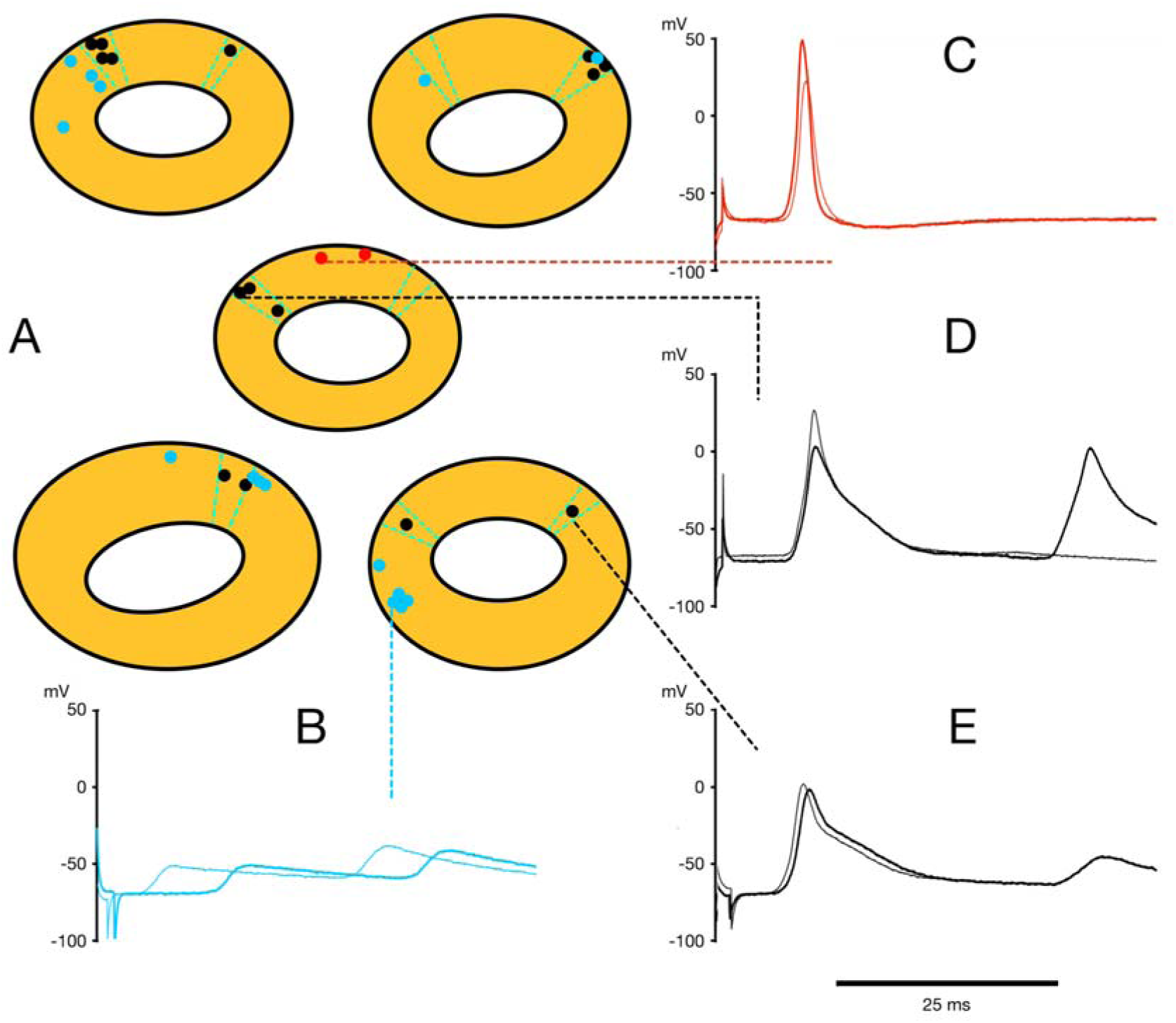
Intracellular recording from identified regions of velum. A) – Chart of velar recording sites traced from five different preparations. The regions of the velum indicated by the dashed green lines show the extent of the visible contraction. Filled circles show recording sites; blue, sites with slow “synaptic” events (B); red, sites with pure action potentials (C); black, sites with a large “synaptic” component and an action potential (sometimes truncated) on the rising phase D, E). B, C, D, E – the thicker lines are typical intracellular records from the indicated recording site (dashed line). The thinner line is a typical response from a near neighbour. Time scale refers to all records. Preparations in filtered seawater plus 10 mM TRIS buffer at pH 7.6. Temperature 10-13°C

The records in Figures 9B-E show the three main kinds of intracellular response to epithelial stimulation. For each pair of records the recording location of the thicker trace is indicated on the chart (Figure 9A), the thinner traces being recorded from neighbouring sites. In 15 sites, marked with filled blue circles, the records took the form of simple synaptic potentials (Figure 9B). In 14 of these 15 sites, the recording location was in a non-contracting area of the velum. Two sites, marked with filled red circles in Figure 9A, were located mid-way between the two sets of Claus’ fibres and records from these sites (Figure 9C) had overshooting action potentials with a distinct undershoot. At another 15 sites, marked with filled black circles in Figure 9A, there was a composite response consisting of a spike-like event arising from the top of a synaptic potential (Figure 9D, E). All 15 of these latter responses were obtained from contracting regions of the velum.

Records from sites with a spike-like component (i.e. Figures 9D, E) had a larger synaptic potential component than those in other regions. This might mean that contracting fibres were located near a major synaptic input although other evidence indicated that inputs are also distributed laterally. In Figure 9B for example the synaptic potential in one record (thinner trace) arises with a significantly shorter delay, the stimulating electrode having been moved from a position near the contracting fibres to one next to the recording site.

One region for which there is no evidence of synaptic input is the sub-delta region (Figure 9C). There is little movement in this area (see Figure 8Cxii) and so any contraction contributes to the overall tension that builds up between the two sets of Claus’ fibres.

#### Synaptic delay

To examine the epithelial inputs to the nerve ring in more detail, isolated nectophores were bisected by cutting between the two sets of Claus’ fibres along the upper and lower nerve tracts. In these experiments the recording site remained constant while the stimulation point ranged widely over the exumbrella epithelium. Figure 10A shows a schematic representation of a divided-nectophore preparation, traced from stored images, with the velum indicated in yellow. The filled black circles show the position of the cactus spines used to stabilize the half-preparation and flatten it as much as possible. The recording site, represented by an open red circle, is located on the velum just outside of the light-sensitive pigmented region (Totton, 1954; Mackie, 1962). The open blue circles represent eight different stimulation sites. The typical extracellular record obtained is shown in Figure 10B (blue trace). The delay between the onset of the stimulus pulse and the initial rise of the response in the velum (green arrowhead) is plotted in Figure 10C (blue symbols) against the distance between stimulus and recording sites. The data points were well fitted by a straight line with a slope showing that the signal travelled with a conduction velocity of 35.7 cm/s. Extrapolation to the y axis showed that the fixed component to the delay was 1.4 ms. This represents the delay in initiating the epithelial impulse plus any synaptic delays experienced by the signal en route to the velum. The regression line R^2^ value is 0.98.

**Figure 10.**
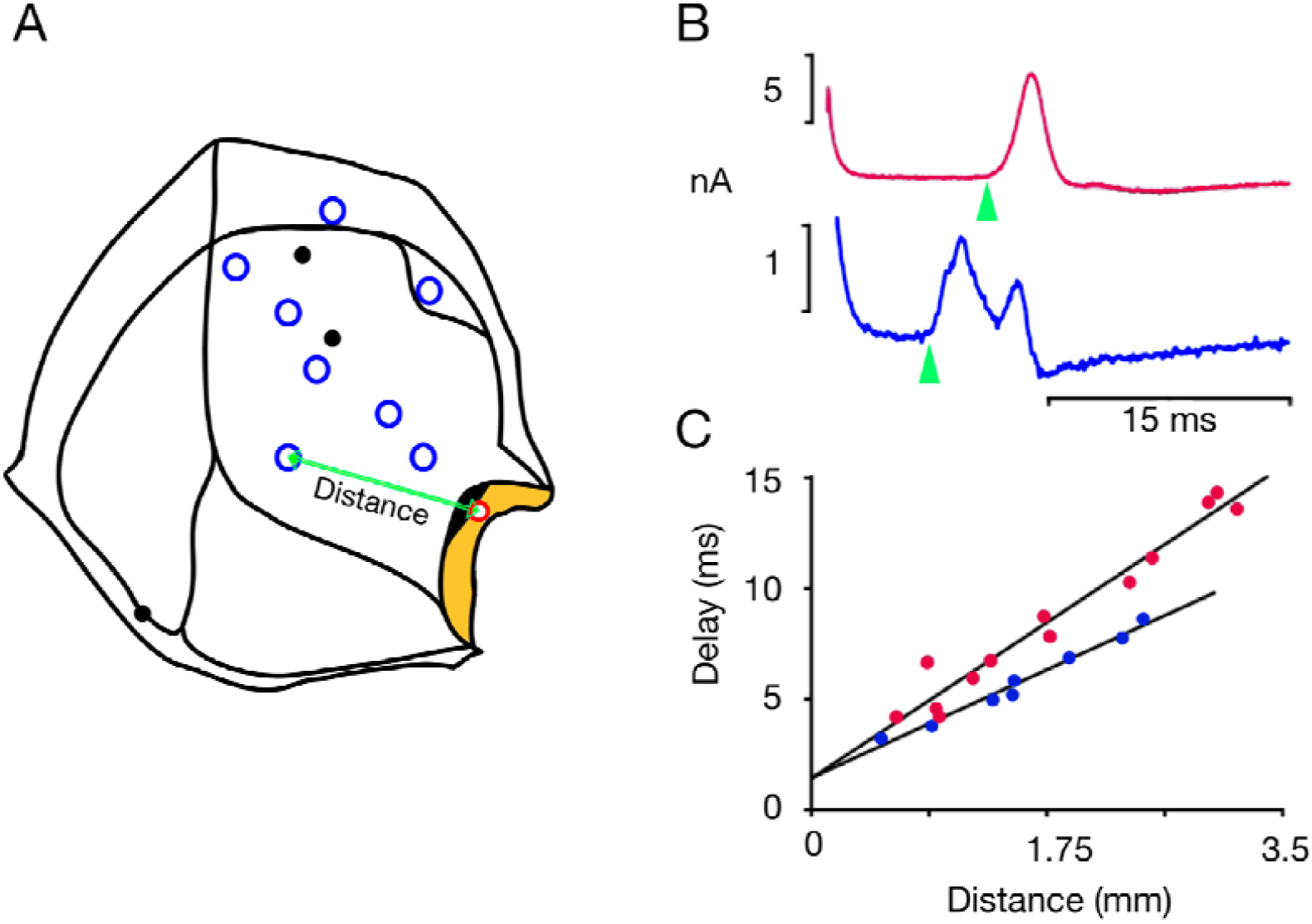
Signal conduction from exumbrella epithelium to the velum. A) – Schematic representation of a half-nectophore preparation pinned flat with cactus spines (filled black circles). The extracellular recording site (open red circle) is located on the surface of the velum (yellow) just outside of the black-pigmented area. The open blue circles show the various positions of the bipolar stimulating electrode. B) – Extracellular recordings of velar muscle responses to electrical stimulation of the exumbrella epithelium; two different preparations. The lower record (blue) is a typical response from the experiment represented in A. The response delay is shown as the time from the onset of the stimulus to the initial rise of the response (arrowhead). C) – delay of the response (ms) plotted against the distance between the stimulating electrode and the recording site (green arrow in A); best fit provided by regression analysis. Preparations in filtered seawater plus 25 mM TRIS buffer at pH 7.6; plus 20% isotonic MgCl_2_ (red data points only). Suction pipette tip diameter, 10-15 μm. Temperature 13-15°C

A similarly high R^2^ value (0.96) was obtained when fitting data from a second experiment (red symbols in Figure 10C). In this case there were twelve different stimulation sites and the velum response was more spike-like (red trace in Figure 10B). The conduction velocity was 24.7 cm/s and the fixed delay was 1.4 ms. Spike-like responses were recorded from three other experiments when the suction pipette was placed at the border between the pigmented area and the Claus’ fibre region. One preparation consisted of an intact nectophore; in the other two the nectophore had been bisected, either as in Figure 10 or at right angles. A similar relationship between conduction time and conduction distance was obtained in each case. The mean value for the conduction velocity was 30 cm/s (± 1.9, SEM; n = 5) and for the delay it was 1.3 ms (± 0.09, SEM; n = 5). The temperature range for these experiments was 11.5-15°C. A trial using a recording site mid-way between the sets of Claus’ fibres and halfway across the velum was unsuccessful because of the variable delay even at a fixed stimulating site. At this site the waveform consisted of a spike preceded by a slow rise to threshold (3 ms ramp) and may be the result of a variable input from multiple remote sites.

Suction electrodes were used to record propagating signals from the exumbrella surface itself (G.O. Mackie personal communication). Three electrodes (one stimulating and two recording) were arranged in a straight line on the surface of an isolated nectophore. When the distance between the recording electrodes was 2.1 mm, the time taken to travel between them was 7.5, and 8 ms in two separate trials (52117 experiment). Thus the conduction velocity was 26-28 cm/s, in line with the values reported here. A synaptic delay of <1.3 ms compares well with the value of 0.7 ms at 12° C in the hydrozoan jellyfish, *Aglantha digitale* (Kerfoot et al., 1985).

To account for the action potentials recorded from the sub-delta region between the two sets of Claus’ fibres (Figure 9C) we assume that current spreading from either side is sufficient to take the membrane to threshold voltage. The question therefore arises as to why current spreading more laterally does not also give rise to overshooting action potentials. The answer to this question has yet to be resolved but one possibility is that the fibres in this region are less well coupled electrically.

## Discussion

If the *Nanomia* colony relies on its nectophores simply for locomotion the nerve networks involved have a surprising degree of complexity. The major network, the nerve ring, is associated with two lateral complexes as well as their associated projections. There is also a “ganglionic” nerve cell cluster in the delta region of the upper nerve tract and, at the stem attachment point, a “terminal group” of cell bodies associated with the lower nerve tract. Added to this, the nectophore’s exumbrella epithelium serves as an adjunct to the nervous system and its propagating impulses are necessary for backward swimming (Mackie, 1964). Here we show that a synaptic link translates epithelial signals into Claus’ muscle excitation. We also discuss the contribution of endodermal muscles to backwards swimming and describe the neuroanatomy of other structures concerned with subsidiary nectophore functions.

### Nematocysts

Although elements of the upper nerve tract overlap individual nematocytes located on the prominent nectophore ridges (Figure 6), they may not make synaptic contact. Similar structures in the siphonophore *Cordalgama cordiformis* are without nerve connections and are thought to be purely defensive (Carre and Carre, 1980). In *Nanomia*, their defensive role may be to entangle debris and stop it getting between individual nectophore bells or from entering them during the refilling process.

### Flask-like structures

Many of the flask-like structures that form clusters at the bell margin have a large opening (Figure 5) as if the cell concerned had released its contents. These supposed gland cells have a good supply of nerves and their location would be ideal for releasing chemical messages or toxins into the water ejected from the nectophore during each swim.

### Stem contact point

Under duress the *Nanomia* colony releases one or more of its nectophores. Once released, the nectophore undergoes a protracted sequence of spontaneous swims, which may serve to distract predators (Mackie et al., 1987). The contact point between the nectophore and the stem is the autotomy site, seen as a pronounced “scar”. On the nectophore side a large muscular pedicel houses an endodermal canal which provides a point of origin for the nectophore’s four radial canals (Mackie, 1964). Next to the “scar” is a small ganglionic structure (Figure 2), made up of the cell bodies of the lower tract axons. In *Hippopodius*, another siphonophore, this is where nervous activity in the stem is translated into epithelial impulses in the nectophore and *vice versa* (Bassot et al., 1978). The translation may involve the facilitation of repeated events and similar processing may take place in *Nanomia*.

### Nectophore swimming

The subumbrella contains no nerve-plexus such as might conduct excitation through the circular swimming muscles. A reasonable conclusion (Mackie, 1960) is that the spread of excitation is myoid in nature, with impulses conducted through the striated muscle sheet itself. Forward swimming requires an intact lower nerve tract (Mackie, 1964), but whether elements in the tract synapse with elements in the outer nerve ring or whether the ring and lower nerve elements are continuous, is not clear (Figure 2B). In either case excitation must pass from the tract to the outer nerve ring and from there to the inner nerve ring and on to the swimming myoepithelium. Electron micrographs show that neurites pass through holes in the mesogloea between the inner and outer nerve rings (Jha and Mackie, 1967).

A natural stimulus for backward swimming occurs when anterior parts of the colony strike some resistant object (Mackie, 1964). Applying an electrical shock can reproduce the effect, reverse swimming being initiated by conducted impulses in the exumbrella epithelium. In isolated nectophores, stimulating any area of the exumbrella surface will evoke Claus’ fibre contraction. Action potentials recorded from the contracting muscles arise from conventional low amplitude, probably synaptic, events (Figure 9). Thus the final input to the Claus’ fibres appears to be a neural one. The site of translation from epithelial to neural propagation is not known. There is a good match between the conduction velocity of the epithelial impulses and the time taken for excitation to travel to the velum and it is likely that the translation step takes place at some lateral site near the nerve ring.

The nerve processes within the exumbrella are discrete units rather than a dense interconnecting network (Figure 3) confirming Mackie’s (1964) methylene blue study. These fine nerve processes, in many cases arising from slightly elongated cell bodies, thread their way between flat epithelial cells while spreading around the upper and back surfaces of each nectophore. They converge on the mid-line and give rise to the upper nerve tract, which enters the nerve ring near its central point. Shorter processes also enter the ring at points away from the central trunk. The upper nerve tract is not necessary for backwards swimming (Mackie, 1964; Figure 10). Its function is unknown although a sensory role seems likely.

### Epithelial conduction

Epithelial conduction provides an effective adjunct to the nervous system in many species but, as with *Nanomia*, links with effectors are generally neural. In the Larvacean *Oikopleura labradoriensis*, where tactile stimulation elicits an escape response, the skin epithelium is connected by gap junctions to axons known to initiate locomotion (Bone and Mackie, 1975). In Hydromedusae such as *Polyorchis* the protective “crumpling” that follows stimulation of the exumbrella epithelium, only occurs if the radial muscles responsible make synaptic contact with neurites in the outer nerve ring (King and Spencer, 1981). In the siphonophore *Hippopodius* the evidence is more indirect. Stimulation of the exumbrellar causes involution of the margin, a response similar to “crumpling” (Bassot et al., 1978). Although an excitable epithelium conducts impulses to the muscles responsible, electron microscopy fails to reveal any synapses or nerves (Bassot et al., 1978). However involution is associated with the inhibition of endogenous swimming, which is hard to account for with a purely electrical model.

### Endodermal muscle

During reverse swimming the ectodermal Claus’ fibres in the velum and the endodermal muscles that abut them at the margin, contract together. Epithelial stimulation causes the Claus’ fibres to shorten but only their distal ends become drawn inwards; there is little distortion at the bell margin (Figure 8B). This absence of movement suggests that the ectodermal and endodermal muscle groups generate equal and opposite forces. At the margin movement is constrained by the isometric contraction of the endodermal fibres, which are anchored to the mesogloea at their distal ends and meet the Claus’ fibres proximally. It is because the Claus’ fibres are less constrained at their distal ends that the velum changes its shape so as to deflect the swimming thrust forwards.

Concerted action involving endodermal and exodermal muscles is highly unusual but does contribute to defensive behaviour in other Hydrozoans (Mackie, 1986). Defensive involution in *Hippopodius* depends on the contraction of ectodermal fibres causing the velum to curl outward while contraction of the endodermal fibres makes the pseudovelum roll inward carrying the curled velum with it (Bossot et al., 1978). Endodermal muscles also contribute to the unusual form of defensive crumpling seen in the limnomedusan *Probosidactyla flavicirrata* (Spencer, 1975). In most hydrozoan species the muscle fibres concerned are in the ectoderm (Mackie and Passano, 1968, *Sarsia, Euphysa*; Mackie and Singla, 1975, *Stomotoca*; King and Spencer, 1981, *Polyorchis*). This difference is notable in view of *Probosidactyla*’s isolated phylogenetic position (Cartwright and Nawrocki, 2010).

### Muscle coordination during backward swimming

During a backwards swim there is: i) a generalized contraction of the circular muscles of the nectophore bell and ii) contraction of the Claus’ fibre/endodermal radial muscle combination. Contraction of the circular muscles and the ectodermal and endodermal radial muscles must be timed to coincide, but how does this coordination come about?

Since reverse swimming occurs even after the lower nerve has been severed all three sets of muscles must be excited directly or indirectly by impulses in the exumbrella epithelium. Either the exumbrella impulse excites a neural input that excites all of them or impulses in the exumbrella spread into the subumbrella at the attachment zone (Mackie 1964).

Coordination of the two groups of radial fibres might come about by: i) independent epithelial inputs, ii) an excitable connection, iii) independent nerve inputs.

i. i) Connections between the ectoderm and endoderm across the mesogloea occur at the margin in *Nanomia* (C. L. Singla; see Bassot et al., 1978) and Mackie (1976) shows a “transmesogloeal tissue bridge”, in which an endodermal process in the stem makes contact with several ectodermal cells. Similar transmesogleal gap junctions occur in *Hydra vulgaris* (Kolenkine and Bonnefoy, 1976; Wood, 1979).
ii. In *Hydra oligactis* gap junctions are seen between an endodermal muscular process and a myoepithelial cell in the ectoderm (Hand & Gobel, 1972). In *Nanomia*, electron micrographs show that “the radial muscle system of the velum …. is connected with an endodermal muscle component through holes in the mesogloea” (Jha and Mackie, 1967; Mackie, 1974), although Figure 8 suggests such connections are rare.
iii. In the Cnidaria nerve tissue can arise in both endoderm and ectoderm (Nakanishi et al., 2012). Electron micrographs show bundles of nerve fibres in the endodermal wall of the ring canal in *Sarsia* (Jha and Mackie, 1967) and in other cnidarian polyps and medusae (Anctil, 2000; Davis, 1974; Grimmelikhuijzen, 1983; Mackie and Singla 1975; Singla 1978; Lin et al., 2001). In *Neoturris, Aglantha* and *Polyorchis* fine endodermal nerves (1 μm diameter), provide an inhibitory input to the swimming pacemaker system during feeding (Mackie and Meech, 2008).

In *Nanomia*, even if the subumbrella swimming muscles are directly excited by epithelial impulses in the exumbrella, the Claus’ fibres require a neural intermediate. The likely place for this is at the junction between the two sets of radial fibres because the nerve ring deviates sharply here, and plunges downward to meet the muscles. It seems probable that the endodermal fibres have a neural input in the same area.

## Acknowledgement

We thank the Director and Staff of Friday Harbor Labs, University of Washington, USA, without whose support this work could not have been completed. We also thank Claudia Mills for her support and endless encouragement. We are indebted to George Mackie for initiating this work and allowing us to include his measurements of epithelial conduction velocity. His insights have been invaluable and one of us in particular (RWM) has benefitted from his patient explanations of siphonophore anatomy. TPN was supported by National Sciences Foundation (NSF) grants 1548121, 1557923, 1645219 and Human Frontier Science Program grant RGP0060/2017.

## Declaration

The authors declare they have no financial or competing interests.

